# Does stomatal patterning in amphistomatous leaves minimize the CO_2_ diffusion path length within leaves?

**DOI:** 10.1101/2023.11.13.566960

**Authors:** Jacob L. Watts, Graham J. Dow, Thomas N. Buckley, Christopher D. Muir

## Abstract

Photosynthesis is co-limited by multiple factors depending on the plant and its environment. These include biochemical rate limitations, internal and external water potentials, temperature, irradiance, and carbon dioxide (CO_2_). Amphistomatous leaves have stomata on both abaxial and adaxial leaf surfaces. This feature is considered an adaptation to alleviate CO_2_ diffusion limitations in productive environments where other factors are not limiting as the diffusion path length from stomate to chloroplast is effectively halved. Plants can also reduce CO_2_ limitations through other aspects of optimal stomatal anatomy: stomatal density, distribution, patterning, and size. A number of studies have demonstrated that stomata are overdispersed on a single leaf surface; however, much less is known about stomatal anatomy in amphistomatous leaves, especially the coordination between leaf surfaces, despite their prevelance in nature and near ubiquity among crop species. Here we use novel spatial statistics based on simulations and photosynthesis modeling to test hypotheses about how amphistomatous plants may optimize CO_2_ limitations in the model angiosperm *Arabidopsis thaliana* grown in different light environments. We find that 1) stomata are overdispersed, but not ideally dispersed, on both leaf surfaces across all light treatments; 2) abaxial and adaxial leaf surface patterning are independent; and 3) the theoretical improvements to photosynthesis from abaxial-adaxial stomatal coordination are miniscule (≪ 1%) across the range of feasible parameter space. However, we also find that 4) stomatal size is correlated with the mesophyll volume that it supplies with CO_2_, suggesting that plants may optimize CO_2_ diffusion limitations through alternative pathways other than ideal, uniform stomatal spacing. We discuss the developmental, physical, and evolutionary constraits which may prohibit plants from reaching the theoretical adaptive peak of uniform stomatal spacing and inter-surface stomatal coordination. These findings contribute to our understanding of variation in the anatomy of amphistomatous leaves.

## Introduction

Stomatal anatomy (e.g. size, density, distribution, and patterning) and movement regulate gas exchange during photosynthesis, namely CO_2_ assimilation and water loss through transpiration. Since waxy cuticles are mostly impermeable to CO_2_ and H_2_O, stomata are the primary entry and exit points through which gas exchange occurs despite making up a small percentage of the leaf area (Lange et al., 1971). Stomata consist of two guard cells which open and close upon changes in turgor pressure or hormonal cues (McAdam and Brodribb, 2016). The stomatal pore leads to an internal space known as the substomatal cavity where gases contact the mesophyll. Once in the mesophyll, CO_2_ diffuses throughout a network of intercellular air space (IAS) and into mesophyll cells where CO_2_ assimilation (*A*) occurs within the chloroplasts (Lee and Gates, 1964). Stomatal conductance and transpiration are determined by numerous environmental and anatomical parameters such as vapor pressure deficit (VPD), irradiance, temperature, wind speed, leaf water potential, IAS geometry, mesophyll cell anatomy, and stomatal anatomy.

Many successful predictions about stomata and other C3 leaf traits can be made by hypothesizing that natural selection should optimize CO_2_ gain per unit of water loss for any given set of environmental parameters, including their variability (Cowan and Farquhar, 1977; Buckley et al., 2017; Sperry et al., 2017). Total stomatal area (size × density) is optimized for operational conductance (*g*_s,op_) rather than maximum conductance (*g*_s,max_) such that stomatal apertures are most responsive to changes in the environment at their operational aperture (Franks et al., 2012; Liu et al., 2021). Stomatal aperture can compensate for maladaptive stomatal densities to an extent (Büssis et al., 2006), but stomatal density and size ultimately determine a leaf’s theoretical *g*_s,max_ (Sack and Buckley, 2016), which is proportional to *g*_s,op_ (Murray et al., 2020). Additionally, low stomatal densities lead to irregular and insufficient CO_2_ supply and reduced photosynthetic efficiency in leaf areas far from stomata (Pieruschka et al., 2006; Morison et al., 2005), while high stomatal densities can reduce water use efficiency (WUE) (Büssis et al., 2006) and incur excessive metabolic costs (Deans et al., 2020). Stomatal density positively co-varies with irradiance during leaf development and negatively co-varies with CO_2_ concentration (Gay and Hurd, 1975; Schoch et al., 1980; Woodward, 1987; Royer, 2001), consistent with optimality predictions. In most species, stomata occur only on the abaxial (usually lower) leaf surface; but amphistomy, the occurrence of stomata on both abaxial and adaxial leaf surfaces, is also prevalent in high light environments with constant or intermittent access to sufficient water (Mott et al., 1982; Jordan et al., 2014; Muir, 2018; Drake et al., 2019; Muir, 2019). Amphistomy effectively halves the CO_2_ diffusion path length and boundary layer resistance by doubling boundary layer conductance (Parkhurst, 1978; Harrison et al., 2020; Mott and Michaelson, 1991). Despite these optimality predictions, stomatal anatomy may be partially constrained by physical and developmental limits on phenotypic expression (Croxdale, 2000; Harrison et al., 2020; Muir et al., 2023).

A number of physical and developmental processes constrain stomatal anatomy traitspace. For example, almost all stomata follow the one cell spacing rule to maintain proper stomatal functioning as guard cell movement requires the rapid exchange of ions with neighboring epidermal cells (i.e. subsidiary cells) (Geisler et al., 2000; Dow et al., 2014). This would prevent stomata from being strongly clustered; however, some species (notably in *Begonia*) appear to benefit from the overlapping vapor shells caused by stomatal clustering in dry environments (Yi Gan et al., 2010; Lehmann and Or, 2015; Papanatsiou et al., 2017). Historically, stomatal patterning in dicot angiosperms was thought to be random with an exclusionary distance surrounding each stomate (Sachs, 1974); however, the developmental controls of stomatal patterning are more complex than random development along the leaf surface. Croxdale (2000) reviews three developmental theories which attempt to explain stomatal patterning in angiosperms: inhibition, cell lineage, and cell cycle, ultimately arguing for a cell cycle based control of stomatal patterning. Pillitteri and Torii (2012) review the short and long distance signalling pathways associated with stomatal spacing and development, which include cell to cell communication and whole plant integration to ensure the proper spacing of stomata across a single leaf surface depending on environmental ques. Much less is known about the development of stomata on the adaxial leaf surface in amphistomatous plants. Stomatal size is additionally constrained by genome size with larger genomes leading to larger minimum guard cell size (Jordan et al., 2015). Despite these limitations, ecophysiological theory still predicts optimal stomatal anatomy, the details of which are discussed below.

The patterning and spacing of stomata on the leaf affects photosynthesis in *C*_3_ leaves by altering the CO_2_ diffusion path length from stomata to sites of carboxylation in the mesophyll. Maximum photosynthetic rate (*A*_max_) in *C*_3_ plants is generally co-limited by biochemistry and diffusion, but modulated by light availability (Parkhurst and Mott, 1990; Manter, 2004; Carriquí et al., 2015). Low light decreases CO_2_ demand by limiting electron transport rate, leading to relatively high internal CO_2_ concentration (*C*_i_) and low *A*_max_ (Kaiser et al., 2016). In contrast, well hydrated leaves with open stomata in high light, photosynthesis is often limited by CO_2_ supply as resistances from the boundary layer, stomatal pore, and mesophyll can result in insufficient CO*C*_2_ supply at the chloroplast to maxmimize photosynthesis (Farquhar et al., 1980; Lehmeier et al., 2017). In this study, we focus primarily on how stomatal patterning affects diffusion, ignoring boundary layer and mesophyll resistances.

To maximize CO_2_ supply from the stomatal pore to chloroplasts, stomata should be uniformly distributed in an equilateral triangular grid on the leaf surface so as to minimize stomatal number and CO_2_ diffusion path length (Parkhurst, 1994). As the diffusion rate of CO_2_ though liquid is approximately 10^4^× slower than CO_2_ diffusion through air, mesophyll resistance is generally thought to be primarily limited by liquid diffusion (Aalto and Juurola, 2002; Evans et al., 2009), but diffusion through the IAS has also been shown to be a rate limiting process because the tortuous, disjunct nature of the IAS can greatly increase diffusion path lengths (Harwood et al., 2021). Additionally, tortuosity is higher in horizontal directions (parallel to leaf surface) than vertical directions (perpendicular to leaf surface) because of the cylindrical shape and vertical arrangement of pallisade mesophyll cells (Earles et al., 2018; Harwood et al., 2021). However, the ratio of lateral to vertical diffusion rate is still largely unknown (Morison et al., 2005; Pieruschka, 2005; Pieruschka et al., 2006). Depending on the thickness of the leaf, porosity of the leaf mesophyll, tortuosity of the IAS, and lateral to vertical diffusion rate ratio, minimizing diffusion path length for CO_2_ via optimally distributed stomata may yield significant increases in CO_2_ supply for photosynthesis and higher *A*_max_.

We hypothesized that natural selection will favor stomatal patterning and distribution to minimize the diffusion path length. In amphistomatous leaves, this would be accomplished by 1) a dispersed, equilateral triangular distribution of stomata on both abaxial and adaxial leaf surfaces and 2) coordinated stomatal spacing on each surface that offsets the position of stomata (Fig. 1). Coordination between leaf surfaces is defined in this study as the occurrence of stomata in areas farthest from stomata on the opposite leaf surface. Additionally, because CO_2_ is more limiting for photosynthesis under high light, we hypothesize that in high light 3) there should be more stomata, and 4) stomata should be more uniformly distributed than in low light. Finally, as stomatal densities are selected for optimal operational aperture, we hypothesize that 5) stomatal length will be positively correlated with the area of the leaf surface to which it is closest. We refer to this as the ‘stomatal zone’, the leaf area surrounding a focal stomate closest to that stomate and therefore the zone it supplies with CO_2_). This way, each stomate can be optimally sized relative to the mesophyll volume it supplies.

**Fig. 1:**
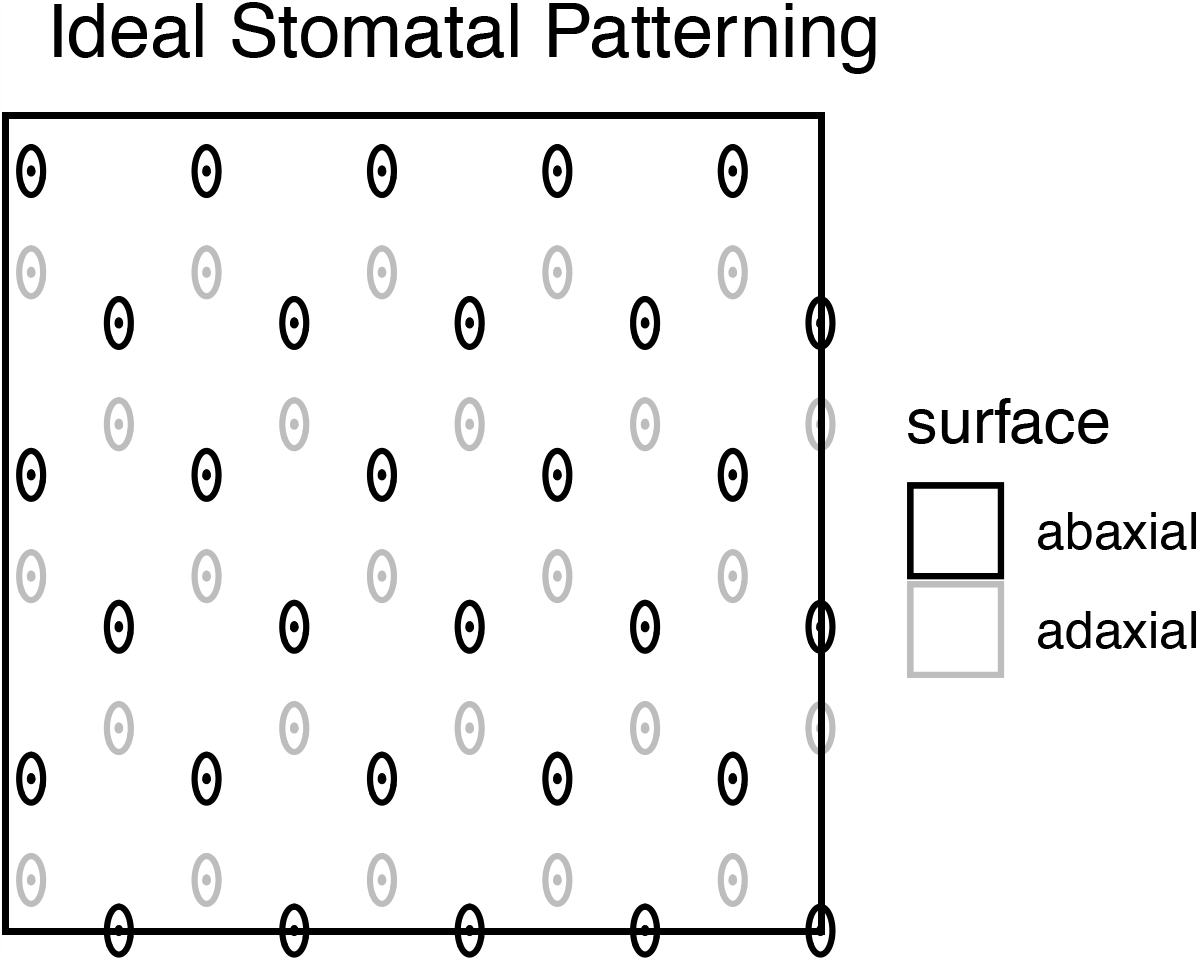
Idealized amphistomatous stomatal grid with uniform stomatal patterning and perfect abaxial-adaxial coordination.

To test these hypotheses, we grew the model plant *Arabidopsis thaliana* in high, medium, and low light and measured stomatal density, size, and patterning on both leaf surfaces, and spatial coordination between them. We use Voronoi tessellation techniques to calculate stomatal zones. We also used a 2-D porous medium approximation of CO_2_ diffusion and photosynthesis to predict the photosynthetic advantage of optimal versus suboptimal coordination in stomatal coordination between surfaces. Specifically, we predicted that traits which affect diffusion path length (leaf thickness, stomatal density, leaf porosity, lateral-vertical diffusion rate ratio), diffusion rate (temperature, pressure), and CO_2_ demand (Rubisco concentration, light) would modulate the advantage of optimal stomatal arrangement following the relationships outlined in Table 1. Here, we integrate over reasonable parameter space to determine the ecophysiological context most likely to favor stomatal spatial coordination in amphistomatous leaves.

**Table 1.**
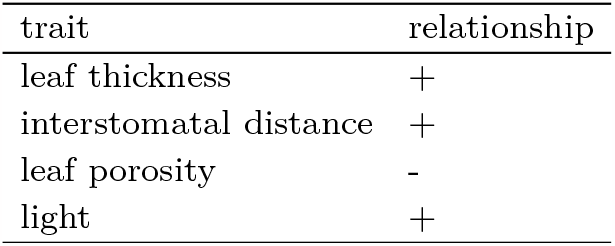
A summary of the hypothesized relationships between leaf traits and environmental conditions and photosynthetic advantage of stomatal spatial coordination in amphistomatous leaves.

| trait | relationship |
| --- | --- |
| leaf thickness | + |
| interstomatal distance | + |
| leaf porosity | - |
| light | + |

## Materials and methods

### Data Preparation

Plant material, growth conditions, and three-dimensional confocal imaging are described in Dow et al. (2017). Briefly, Columbia (Col-0) ecotype of *Arabidopsis thaliana* (L.) Heynh. plants were grown in three different light environments: low light (PAR = 50 *µ*mol m^−2^ s^−1^), medium light (100 *µ*mol m^−2^ s^−1^), and high light (200 *µ*mol m^−2^ s^−1^). PAR stands for photosynthetically active radiation. Seeds were surface-sterilized and stratified at 4°C for 3–5 d in 0.15% agarose solution and then sown directly into Pro-Mix HP soil (Premier Horticulture; Quakerstown, PA, USA) and supplemented with Scott’s Osmocote Classic 14-14-14 fertilizer (Scotts-Sierra, Marysville, OH, USA). At 10–14 d, seedlings were thinned so only one seedling per container remained. Plants were grown to maturity in growth chambers where the conditions were as follows: 16 : 8 h, 22 : 20°C, day : night cycle. Imaging of the epidermis and internal leaf structures was performed using a Leica SP5 confocal microscope with the protocol developed by Wuyts et al. (2010) with additional modification described in Dow et al. (2017). We captured 132 images in total, making 66 abaxial-adaxial image pairs. We measured stomatal position and size using ImageJ (Schneider et al., 2012).

### Single surface analyses

We tested whether stomata are non-randomly distributed by comparing the observed stomatal patterning to a random uniform pattern. For each leaf surface image with *n* stomata we generated 10^3^ synthetic surfaces with *n* stomata uniformly randomly distributed on the surface. For each sample image, we compared the observed Nearest Neighbor Index (NNI) to the null distribution of NNI values calculated from the synthetic data set. NNI is the ratio of observed mean distance 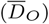 to the expected mean distance 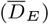 where 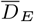 is:

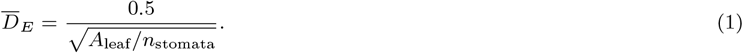

*A*_leaf_ is leaf area visible in the sampled field and *n*_stomata_ the number of stomata. 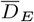 is the theoretical average distance to the nearest neighbor of each stomate if stomata were uniformly randomly distributed (Clark and Evans, 1954). 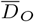 calculated for each synthetic data set is:

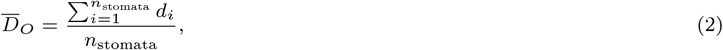

where *d*_*i*_ is the distance between stomate_*i*_ and its nearest neighbor. We calculated NNI using the *R* package **spatialEco** version 2.0.1 (Evans and Murphy, 2023). The observed stomatal distribution is dispersed relative to a uniform random distribution if the observed *NNI* is greater than 95% of the synthetic NNI values (one-tailed test). We adjusted *P* -values to account for multiple comparisons using the Benjamini-Hochberg (Benjamini and Hochberg, 1995) false discovery rate procedure implemented in the *R* package **multtest** version 2.56.0 (Pollard et al., 2005).

For each sample image, we also simulated 10^3^ synthetic data with *n* stomata ideally dispersed in an equilateral triangular grid. For these grids, we integrated over plausible stomatal densities and then conditioned on stomatal grids with exactly *n* stomata. The simulated stomatal count was drawn from a Poisson distribution with the mean parameter *λ* drawn from a Gamma distribution with shape *n* and scale 1 (*λ* ∼ Γ(*n*, 1)). Γ(*n*, 1) is the posterior distribution of *λ* with a flat prior distribution. This allows us to integrate over uncertainty in the stomatal density from the sample image.

We developed a dispersion index DI to quantify how close observed stomatal distributions are to random uniform versus maximally dispered in an equilateral triangular grid. DI varies from zero to one, where zero is uniformly random and one is ideally dispersed:

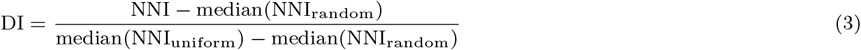

NNI is calculated for each sample image as described above; median(NNI_random_) and median(NNI_uniform_) are calculated from the synthetic data specific to each sample image as described above. We tested whether light treatment affects DI and stomatal density (*D*_*S*_) using analysis of variance (ANOVA).

Finally, we examined the relationship between stomatal zone area and stomatal length using a Bayesian linear mixed-effects model fit with the *R* package **brms** version 2.20.4 (Bürkner, 2017, 2018) and *Stan* version 2.33.1 (Stan Development Team, 2023). Stomatal zone area was calculated using Voronoi tessellation (e.g. Fig. 2). The stomatal zone area, *S*_area_, is the region of the leaf surface whose distance to stomate, *S*, is less than the distance to any other stomate, *S*. Stomatal length was measured in ImageJ (Schneider et al., 2012). We modeled fixed effects of surface, light treatment, stomatal length, and their 2- and 3-way interactions on 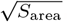. We included random intercepts, random effects of surface, random slopes, and random surface-by-slope interactions within both plant and individual to account for nonindependence of stomata within the same plant or individual. We also modeled residual variance as a function of light treatment. We sampled the posterior distribution from 4 chains with 1000 iterations each after 1000 warmup iterations. We calculated convergence diagnostics 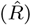 and effective sample sizes following Vehtari et al. (2021). We estimated the marginal slope and 95% highest posterior density (HPD) intervals between stomatal length and 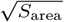 using the *emtrends* function in the *R* package **emmeans** version 1.8.8 (Lenth, 2023).

**Fig. 2:**
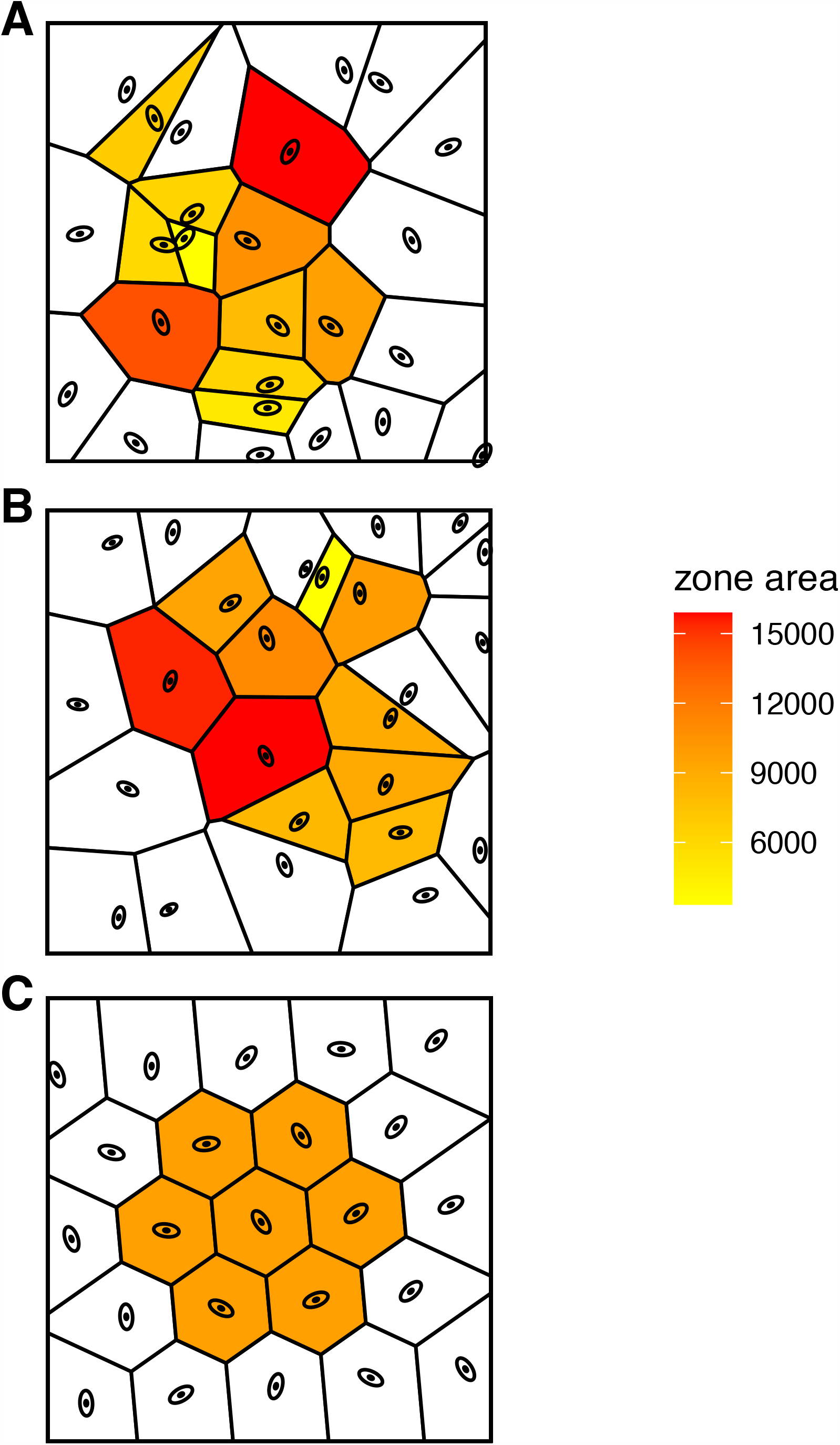
Examples of synthetic and real leaf surfaces. A) Uniform random synthetic leaf surface; B) Example of real leaf surface; C) Regularly distributed synthetic leaf surface. The zone defined by each stomate was calculated with voronoi tessellation and correlated with stomatal length in real leaves.

### Paired Abaxial and Adaxial Surface Analysis

To test whether the position of ab- and adaxial stomata are coordinated we compared the observed distribution to a null distribution where the positions on each surface are random. For each pair of surfaces (observed or synthetic) we calculated the distance squared between each pixel to the nearest stomatal centroid with the *R* package **raster** version 3.6.26. We refer to this as the ‘nearest stomatal distance’ or NSD. Then we calculated the pixel-wise Pearson correlation coefficient. If stomatal positions on each surface are coordinated to minimize the distance between mesophyll and the nearest stomate, then we expect a negative correlation. A pixel that is far from a stomate on one surface should be near a stomate on the other surface (Fig. 1). We generated a null distribution of the correlation coefficient by simulating 10^3^ synthetic data sets for each observed pair. For each synthetic data set, we simulated stomatal position using a random uniform distribution, as described above, matching the number of stomata on abaxial and adaxial leaf surfaces. Stomatal positions on each surface are coordinated if the correlation coefficient is greater than 95% of the synthetic correlation values (one-tailed test).

### Modeling Photosynthesis

We modeled photosynthesis CO_2_ assimilation rate using a spatially-explicit two-dimensional reaction diffusion model using a porous medium approximation (Parkhurst, 1994) using the finite element method (FEM) following Earles et al. (2017). Consider a two-dimensional leaf where stomata occur on each surface in a regular sequence with interstomatal distance *U*. The main outcome we assessed is the advantage of offsetting the position of stomata on each surface compared to have stomata on the same *x* position on each surface. With these assumptions, by symmetry, we only need to model two stomata, one abaxial and one adaxial, from *x* = 0 to *x* = *U/*2 and from the adaxial surface at *y* = 0 to the abaxial surface at *y* = *L*, the leaf thickness. We arbitrarily set the adaxial stomate at *x* = 0 and toggled the abaxial stomata position between *x* = *U/*2 (offset) or *x* = 0 (below adaxial stomate). The ‘coordination advantage’ of offset stomatal position on each surface is the photosynthetic rate of the leaf with offset stomata compared to that with stomata aligned in the same *x* position:

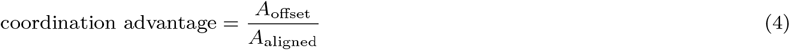

We modeled the coordination advantage over a range of leaf thicknesses, stomatal densities, photosynthetic capacities, and light environments to understand when offsetting stomatal position on each surface might deliver a significant photosynthetic advantage (Table 2). The complete model description is available in the Supporting Information.

**Table 2.**
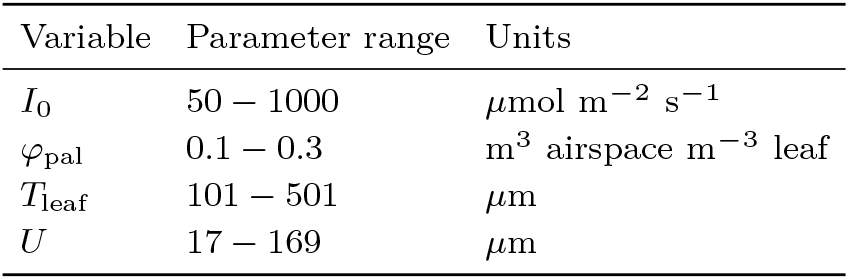
The parameter range of model variables tested for their effect on coordination advantage (Equation 4) using a 2-D porous medium approximation. We used regularly spaced values within each range and simulated across all combinations. Here we converted model units to more conventional units (e.g. m to *µ*m). *I*_0_: PPFD incident on the leaf surface; *ϕ*_pal_: Fraction of intercellular airspace (aka porosity), palisade; *T*_leaf_: Leaf thickness; *U* : Interstomatal distance.

## Results

Stomatal density of *Arabidopsis thaliana* varies among light treatments (ANOVA, *F*_2,126_ = 681,*P* = 2.88 × 10^−68^) because the density is much greater in the high light treatment (Fig. 3). Density is consistently greater on abaxial leaf surfaces across all light treatments (ANOVA, *F*_1,126_ = 44.2,*P* = 8.21 ×10^−10^; Fig. 3). There is no evidence for an interaction between light treatment and surface (ANOVA, *F*_2,126_ = 2.75 × 10^−2^,*P* = 0.973). Leaves are amphistomatous with a mean stomatal density ratio of 0.44.

**Fig. 3:**
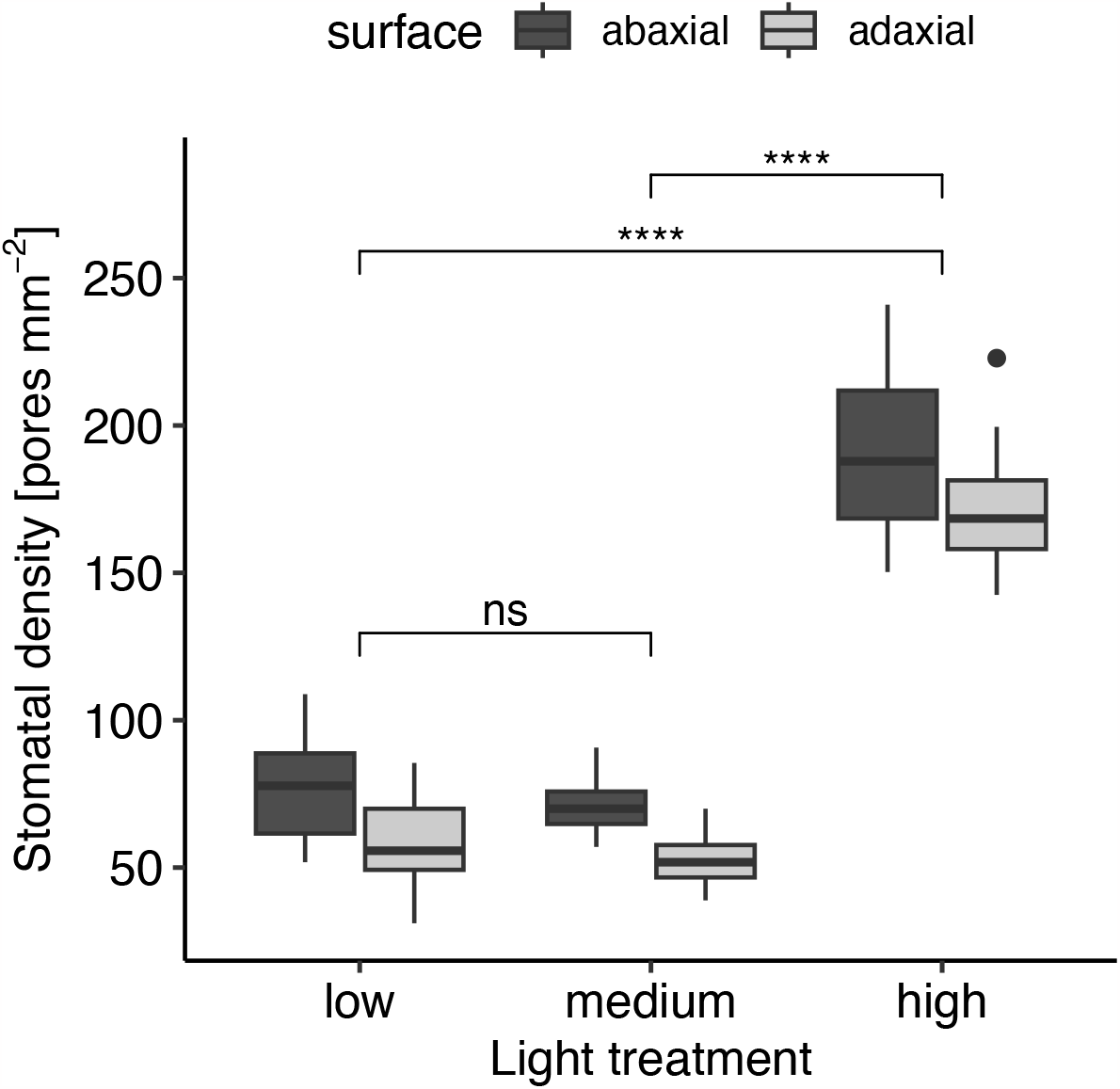
Stomatal density is higher in *A. thaliana* plants grown under high light conditions. We determined statistical significance between light treatments using Tukey post-hoc tests. ^*^ 0.05 > *P* ≥ 0.01;^**^ 0.01 > *P* ≥ 0.001;^***^ 0.0001 > *P* ≥ 0.0001;^***^ *P* < 0.0001.

### Stomatal distribution is non-random, but far from ideal

Many leaf surfaces (37 of 132, 28%) are significantly overdispersed compared to a random uniform distribution, but none were close to an ideal hexagonal pattern (dispersion index = 1; Fig. 4). Before controlling for multiple comparisons, 43.2% are significantly overdispersed. The dispersion index differs significantly among light treatments (ANOVA, *F*_2,126_ = 8.55,*P* = 3.30 × 10^−4^) because the medium light treatment is significantly less than the low treatment (Fig. 4). Dispersion index is consistently greater on adaxial leaf surfaces across all light treatments (ANOVA, *F*_1,126_ = 28.8,*P* = 3.67 × 10^−7^; Fig. 4). There is no evidence for an interaction between light treatment and surface (ANOVA, *F*_2,126_ = 0.577,*P* = 0.563).

**Fig. 4:**
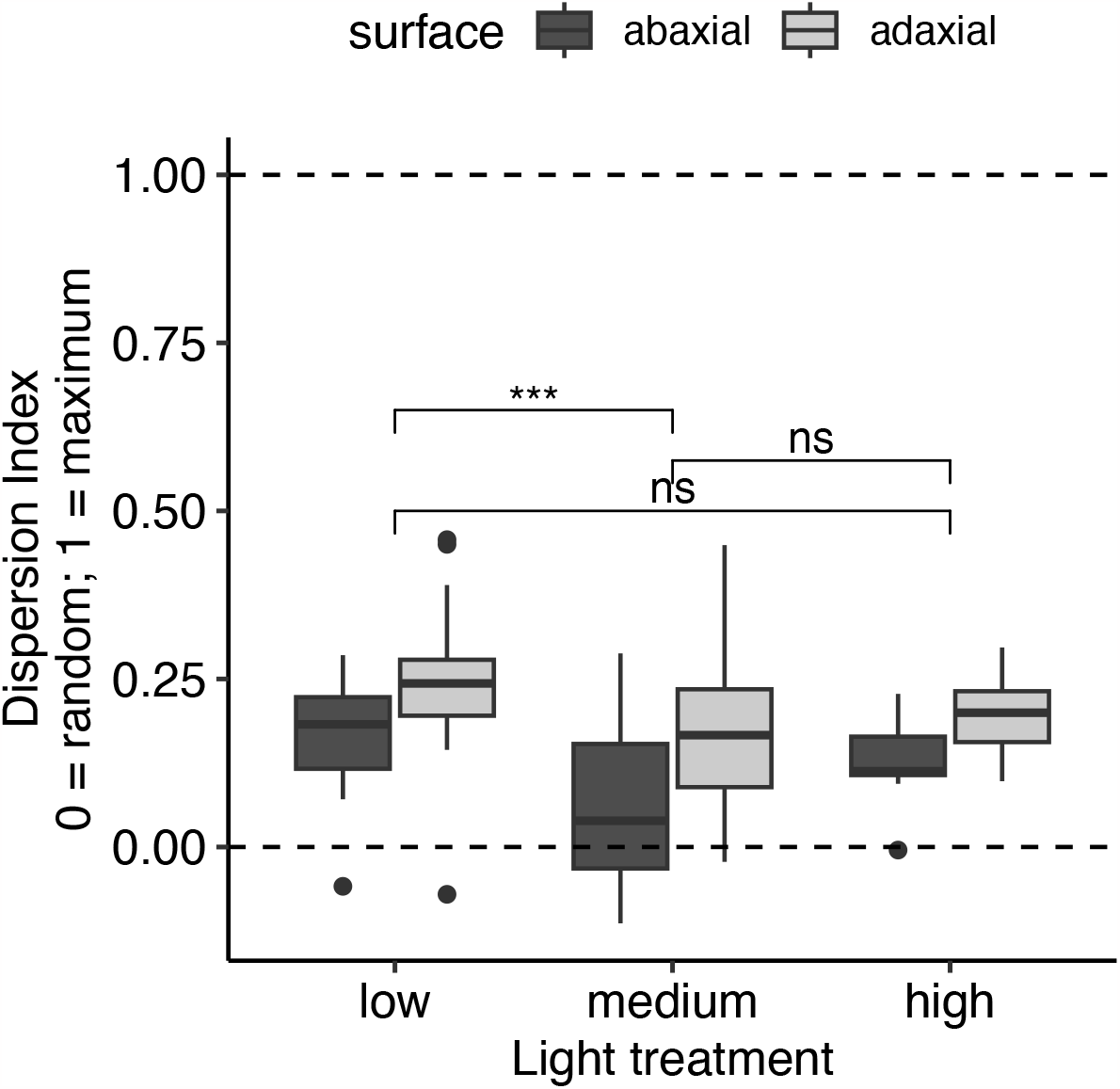
Stomata are more dispersed than expected under the null model of uniform random position (dispersion index = 0) but far from a distribution that maximizes distance between stomata (dispersion index = 1). We determined statistical significance between light treatments using Tukey post-hoc tests. ^*^ 0.05 > *P* ≥ 0.01;^**^ 0.01 > *P* ≥ 0.001;^***^ 0.0001 > *P* ≥ 0.0001;^***^ *P* < 0.0001.

### No evidence for coordinated stomatal position between surfaces

There is no evidence of spatial coordination between abaxial and adaxial leaf surfaces. The pixel-wise correlation between nearest stomatal distance (NSD) squared on paired abaxial and adaxial leaf surfaces is not significantly less than zero in any of the 66 leaves (Fig. 5). Before controlling for multiple comparisons, 3% are significantly *positively* correlated. The NSD correlation is not different among light treatments (ANOVA, *F*_2,63_ = 2.28,*P* = 0.111; Fig. 5).

**Fig. 5:**
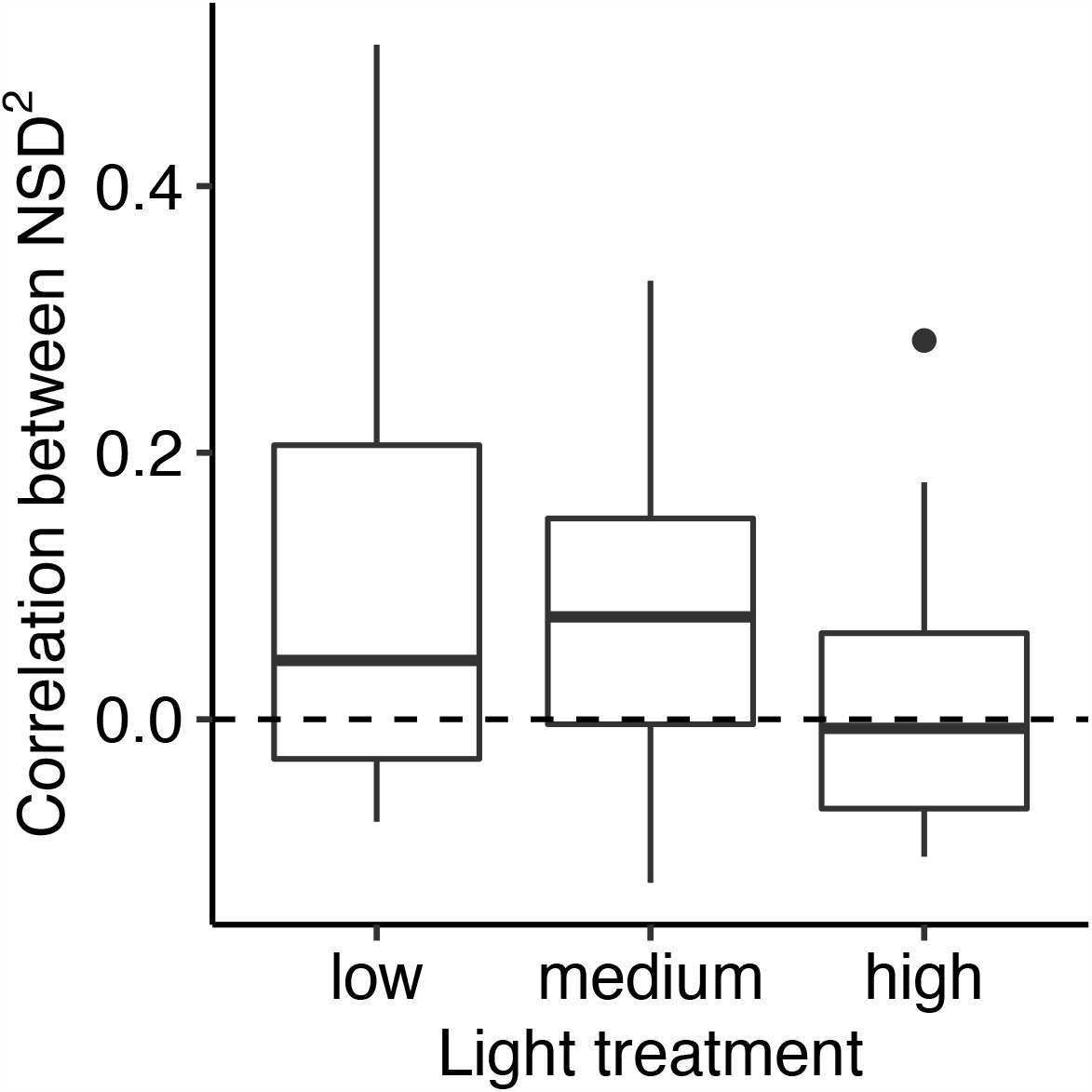
Pixel-wise correlation between near stomatal distance (NSD) squared on paired abaxial and adaxial leaf surfaces. Dashed line indicates zero correlation. Weak positive correlations are not significantly different from zero after correcting for multiple comparisons. The correlation does not differ among light treatments.

### Larger stomata supply larger mesophyll volumes

All parameters converged 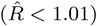 and effective sample sizes were exceeded 10^3^. Across all light treatments and leaf surfaces, stomatal length and stomatal area are weakly positively correlated (Fig. 6). The slope was significantly greater than zero for all abaxial surfaces, but not for the adaxial surface in low and medium light treatments. The estimated marginal slopes and 95% HPD intervals for each combination of light and surface is: low light, abaxial surface: 1.928 [0.779–3.133]; low light, adaxial surface: 1.745 [−0.041–3.373]; medium light, abaxial surface: 1.085 [0.328–1.957]; medium light, adaxial surface: 0.656 [−0.399–1.691]; high light, abaxial surface: 0.597 [0.316–0.911]; high light, adaxial surface: 1.269 [0.831–1.721].

**Fig. 6:**
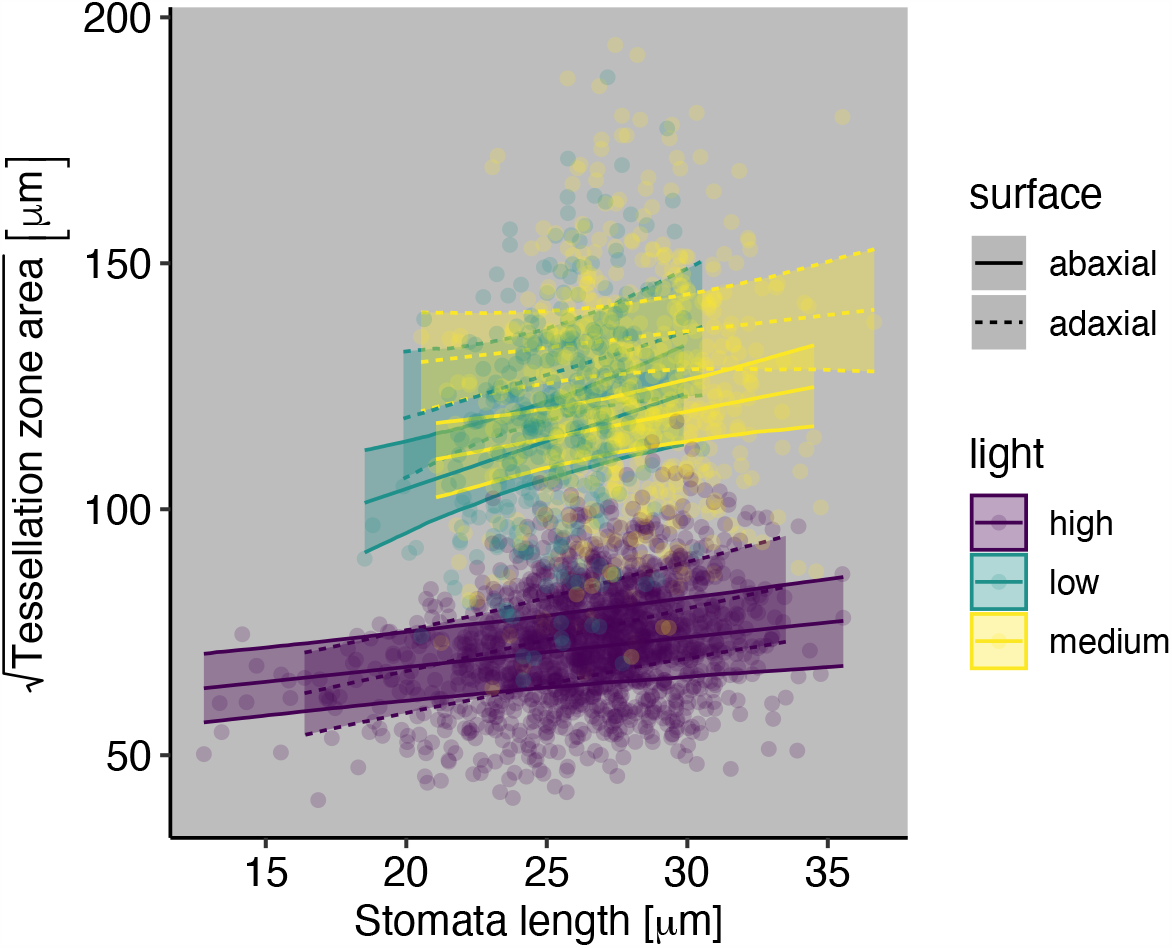
Stomatal length and stomatal zone area are positively correlated. Linear regression lines and 95% confidence ribbons are from a Bayesian linear mixed-effects model.

### Little benefit of coordinated stomatal arrangement

We used the finite element method (FEM) to model CO_2_ diffusion within the leaf and photosynthesis as a 2-D porous medium. Across all realistic parts of parameter space, the coordination advantage is much less than 0.01 (Fig. 7). For reference, a log-response of ratio is 0.01 is approximately 1%. The only exception was for thin leaves (*T*_leaf_ = 100 *µ*m) with few stomata (*U* = 338 *µ*m, which corresponds to a stomatal density of ≈ 10 mm^−2^), where lateral diffusion is major constraint on CO_2_ supply. However, such thin leaves with so few stomata are uncommon among C_3_ plants (some CAM plants have low stomatal density (Males and Griffiths, 2017)). In other areas of parameter space, lateral diffusion limitations were small relative to those along the ab-adaxial axis (see Fig. S1 for a representative model solution).

**Fig. 7:**
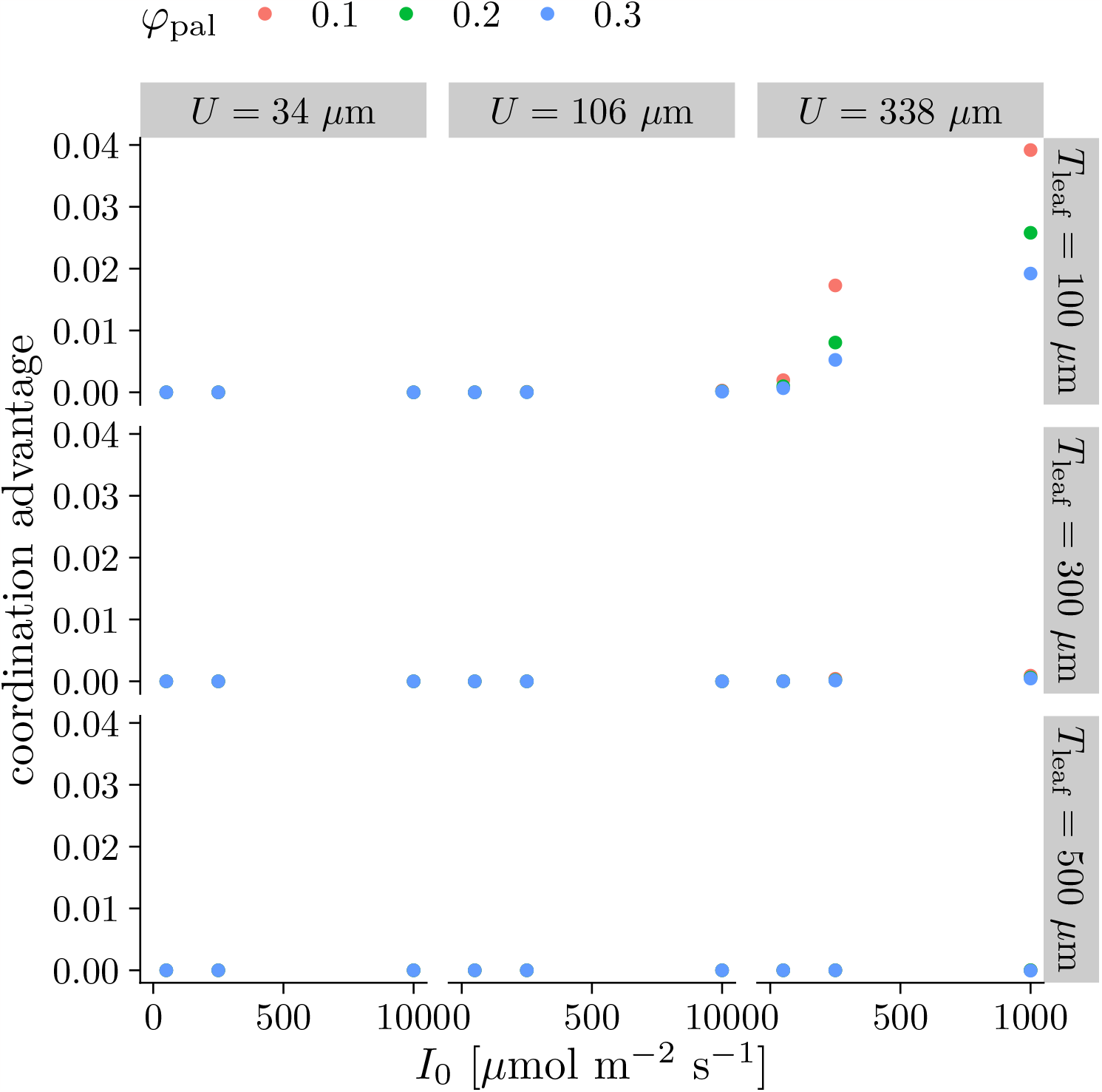
There is little photosynthetic benefit of offsetting stomatal position each surface based on a 2-D model of photosynthesis. The coordination advantage (Equation 4) is close to zero under nearly all of the parameter space Table 2, meaning that the photosynthetic rate of amphistomatous leaves with stomata optimally offset is nearly equal to leaves with stomata on each surface in the same position along the leaf plane. *I*_0_: PPFD incident on the leaf surface; *ϕ*_pal_: Fraction of intercellular airspace (aka porosity), palisade; *T*_leaf_: Leaf thickness; *U* : Interstomatal distance.

## Discussion

Stomata cost resources to maintain (Deans et al., 2020) and expose leaves to risks such as hydraulic failure (Wang et al., 2020) or infection by plant pathogens (Melotto et al., 2017). Therefore leaves should develop enough stomata to adequately supply CO_2_ to chloroplasts, but not overinvest. A widespread hypothesis in plant ecophysiology is that natural selection optimizes traits like stomatal size, density, and distribution to maximize carbon gain relative to any costs in a given environmental context. In principle, spacing stomata to minimize the average distance between stomatal pores and chloroplasts within the mesophyll should increase carbon gain, all else being equal. However, reducing this distance to its absolute minimum may be constrained by developmental processes, or the photosynthetic benefit may be too small to be ‘seen’ by natural selection (i.e. the selection coefficient is less than drift barrier *sensu* Sung et al. (2012)).

We tested five related hypotheses about stomatal spacing in amphistomatous leaves using the model angiosperm *Arabidopsis thaliana* grown under different light intensities. First, we predicted that stomata on each surface are overdispersed relative to a random uniform distribution, which should increase CO_2_ supply. Stomata on each surface are overdispersed (Fig. 4), but are not ideally dispersed in an equilateral triangular grid as would be optimal to minimize CO_2_ diffusion path length and equalize the area supplied by each stomate (Fig. 2). Second, we predicted that an optimal amphistomatous leaf has offset stomata such that stomata are more likely to appear on one leaf surface if there is not a stomata directly opposite it on the other surface as shown in Fig. 1. However, there is no evidence for coordination and the positions on each surface appear independent, regardless of light treatment (Fig. 5). Third, we predicted that plants respond plastically to higher light intensity by increasing stomatal density. *Arabidopsis* plants grown under high light had higher stomatal density than the same genotype growns under low and medium light intensity (Fig. 3). However, we found no support for our fourth prediction that stomata would be more evenly dispersed at high light intensity (Fig. 4). Finally, we predicted that within leaf variation in stomatal size would correlate with stomatal spacing, as larger stomata can supply larger volumes of adjacent mesophyll. In all three light treatments, stomatal size positively covaried with the stomatal zone, i.e. adjacent region of mesophyll that would be supplied by that stomate (Fig. 6).

Stomatal spacing on *A. thaliana* leaves partially supports our overall hypothesis that natural selection minimizes the average distance between stomata and chloroplasts, for a given overall stomatal density. There are three non-mutually exclusive hypotheses for why several of our predictions were wrong. First, our predictions are wrong becuase they are based on overly simplistic assumptions about epidermal and mesophyll anatomy. Second, natural selection may be constrained by developmental processes that prevent phenotypes from reaching their adaptive optima. Third, the benefit of some traits may be of too little consequence to result in fitness differences large enough to respond to selection. We consider the plausibility of these alternative hypotheses below and present ideas for future work to test them.

We assume an idealized leaf epidermal and mesophyll structure that is homogenous and unconstrained by other tradeoffs. Real leaves not only provide pathways for CO_2_ diffusion, but must supply water, intercept light, and deter herbivores and pathogens. These competing interests result in nonuniform epidermal and mesophyll structure that could alter our predictions about optimal stomatal spacing. In order to maintain consistent leaf water potential across the lamina, stomatal density must be coordinated with vein density (Fiorin et al., 2016). Thus, stomatal spacing may be optimized not at the interstomatal level, but at a higher level, coordinating water transport and water loss. We also assume a uniform mesophyll; however, in ab-adaxially differentiated leaves, this assumption is overly simplistic. The pallisade mesophyll is more tightly packed than the spongy mesophyll as an adaptation to intercept light efficiently, so lateral diffusion may be more limiting in the adaxial portion of the leaf. This may explain why adaxial leaf surfaces have consistently higher dispersion indices than abaxial surfaces across all light treatments (Fig. 4). Future gas exchange models should include multiple parameter sets for different strata of mesophyll tissue.

No developmental pathway exists to ensure an idealized placement of stomata on the leaf surface. Rather, stomatal development is a dynamic process that must be plastic to environmental cues. Leaves develop based on short and long distance signalling pathways which relay information about incoming light, humidity, temperature, and surrounding stomata to developing leaf tissues (Pillitteri and Torii, 2012). Our results showing an intermediate level of dispersion in stomatal spacing may be best explained by these developmental pathways which ensure the proper spacing of stomata, with an added random effect brought about by the necessity for plasticity in stomatal development (Fig. 4). However, any nonideal randomness in stomatal spacing may offset by simultaneous and coordinated development of the IAS (Baillie and Fleming, 2020). The fact that stomata which supply a greater mesophyll volume tend to be larger suggests that plants may use coordinated development of multiple leaf tissues to negate nonideal stomatal spacing (Fig. 6).

In amphistomatous leaves, ideal stomatal spacing is complicated by a third dimension. Our gas exchange model demonstrates little photosynthetic gain from abaxial-adaxial stomatal coordination (Fig. 7). Even though lateral diffusion may limit photosynthesis (Morison et al., 2005), the marginal gain from optimally offsetting stomata is not sufficient to generate fitness differences relative to the strength of genetic drift (i.e. the drift-barrier). We can similarly extrapolate that an ideal, equilateral triangular stomatal spacing is only slightly better than a suboptimal pattern. Any benefit garnered by ideal stomatal spacing may be additionally offset by a cost to developmental flexibility in variable environments (Pillitteri and Torii, 2012; Baillie and Fleming, 2020). Explaining these observations as the result of weak selection is in tension with the finding that stomatal size and zone positively covary, which would suggest that small changes in lateral diffusion distance are significant. As described above, the positive correlation between stomatal size and zone may be explained by common developmental processes rather than as an adaptation to maximize CO_2_ diffusion. In any case, there is no evidence for coordinated development of both leaf surfaces, and very little theoretical benefit to photosynthesis, except in marginal circumstances which are exceptionally rare in nature.

Our study corroborates previous studies which demonstrate that stomata are non-randomly distributed along the leaf surface as a result of developmental mechanisms such as spatially biased arrest of stomatal initials (Boetsch et al., 1995), oriented asymmetric cell division (Geisler et al., 2000), and cell cycle controls (Croxdale, 2000). We do not investigate the potential developmental pathways that influence stomatal dispersion in this study; however, they are important to consider as these pathways could limit plants from reaching a theoretical peak in the adaptive landscape: uniform stomatal dispersion. Instead, as this study suggests, plants may simply compensate with higher stomatal density and by fitting stomatal size to the area that they supply with CO_2_. To understand why stomata are not ideally dispersed, more modeling (with a more complex set of assumptions including vein density and IAS structure) should be done to estimate the fitness gain of stomatal dispersion. Additionally, genetic manipulation studies should attempt to create mutants with clustered and ideally dispersed stomata for a comparison of their photosynthetic traits. This could have extremely important implications for maximum assimilation rates in crops as most crop species are grown in high light where CO_2_ is often limiting. In drought-prone environments, increased stomatal dispersion may increase water use efficiency by reducing the number of stomata needed to achieve the same internal CO_2_ concentration, *C*_i_.

Our results suggest that after optimizing stomatal density and having developmental rules for spacing stomata relatively evenly, there may be limited gains to further optimization. Therefore, developmental constraints may be necessary to make sense of some features of stomatal spacing and distribution. The possibility that ideal stomatal spacing is not the “tallest” fitness peak must also be explored, as stomate size is demonstrated in this study to covary with mesophyll volume supplied with CO_2_. This may be especially true in highly variable environments or in large tree species with sun and shade leaves where developmental cues may change rapidly. The temporal component, not considered here, could also have significant implications, as CO_2_ may only be limiting to photosynthesis during short periods when all other conditions are ideal. In these cases, the theoretical benefits of ideal stomatal spacing are further diminished. Future exploration of these competing hypotheses would require more advanced modeling, additional exploration of IAS space development and its effects on gas exchange, both real and modeled, and knowledge about how often the species of interest is CO_2_ limited across of range of natural settings. Despite these additional considerations, this study represents an important contribution to understanding the potential drivers of and limitations to stomatal anatomy in amphistomatous plants.

## Author contributions statement

JLW and CDM conceived of the project, analyzed data, and wrote the manuscript. GJD provided data. TNB contributed to model development and helped edit the manuscript.

