## Supplementary Information for "Does stomatal patterning in amphistomatous leaves minimize the CO_2_ diffusion path length within leaves?"

### 2-D Photosynthesis Model

We modeled leaf photosynthesis using a two-dimensional porous medium approximation. The `steady.2d()` function in the *R* package **rootSolve** version 1.8.2.4 (Soetaert and Herman 2009) solves the model using a finite element method (FEM). The 2-D leaf profile is $n_x$ elements long and $n_z$ elements deep with square elements of area $t_{\text{elem}}^2$. In all cases, we set $t_{\text{elem}} = 1\ \mu\text{m}$. Table S1 is a glossary model terms and symbols.

##### Leaf anatomy

We assume that the leaf is a homogenous 2-D medium. The mesophyll is $T_{\text{leaf}}$ thick and the stomata are regularly spaced apart by distance $U$ on both ab- and adaxial surfaces. In this scenario, we assume that the stomata on each surface are precisely offset from each other by distance $U/2$. This minimizes the average distance between any point in the mesophyll and its nearest stomate. Because of the regular spacing, we only need to model the region between a stomate on surface and the next stomate on the other surface (Fig. S1). The rest of the mesophyll will be the same because of symmetry. This allowed us to set the boundary fluxes on the left and right sides of the leaf profile to 0.

##### Solving within-leaf gradients in $CO_2$ assimilation and concentration

We extended the 1-D FEM of Earles et al. (2017) to solve a set of partial differential equations describing $CO_2$ diffusion, photosynthesis, and respiration throughout a 2-D leaf geometry. The diffusive flux of $CO_2$ through ab- and adaxial stoamta, intercellular airspace, and mesophyll cells in the $x$ (length) and $z$ (depth) dimensions is:

$$D_{\text{e}} \nabla^2 C_{\text{ias}} = D_{\text{e}} \left( \frac{\partial^2 C_{\text{ias}}}{\partial x^2} + \frac{\partial^2 C_{\text{ias}}}{\partial z^2} \right) = -f_{\text{liq}}, \tag{S1}$$

where

$$f_{\text{liq}} = r_{\text{d}} + r_{\text{p}} - r_{\text{c}} \tag{S2}$$

and

$$D_e = \frac{\varphi}{\tau} D_c \quad \text{(S3)}$$

is the effective diffusivity of $CO_2$ though a porous medium composed of an intercellular airspace with a porosity ($\varphi$; $m^3$ airspace $m^{-3}$ leaf) and tortuosity ($\tau$; m $m^{-1}$). We assume the palisade ($\varphi_{pal}$) is less porous than the spongy ($\varphi_{spg}$) mesophyll (Table S1). $D_c$ is the diffusion coefficient (m $s^{-1}$) for $CO_2$ in the intercellular airspace, $C_{ias}$ is the [$CO_2$] (mol $m^{-3}$) at horizontal positions $x$ and depth $z$ in the intercellular airspace, $f_{liq}$ is the volumetric rate of $CO_2$ diffusion from the intercellular airspace into the chloroplast stroma (mol $m^{-3}$ $s^{-1}$), $r_c$ is the volumetric rate of ribulose 1,5-bisphosphate (RuBP) carboxylation (mol $m^{-3}$ $s^{-1}$), $r_d$ is the volumetric respiration rate (mol $m^{-3}$ $s^{-1}$), and $r_p$ is the volumetric photorespiration rate by Rubisco (mol $m^{-3}$ $s^{-1}$). Following Earles et al. (2017), $r_d$ is assumed constant per stroma surface area (Table S1) and $r_p$ is a function of carboxylation ($r_c$) and $C_{liq}$:

$$r_p = r_c \frac{\Gamma^*}{C_{liq}}. \quad \text{(S4)}$$

Carboxylation rate is the minimum of the Rubisco-limited ($w_c$) or RuBP-regeneration limited ($w_j$) carboxylation rates:

$$r_c = \min(w_c, w_j) \frac{S_m}{T_{leaf}} V_{strom}, \quad \text{(S5)}$$

where

$$w_c = \frac{k_c X_c C_{liq}}{K_m + C_{liq}}, \text{ and} \quad \text{(S6)}$$

$$w_j = \frac{C_{liq} j_e}{4C_{liq} + 8\Gamma^*}. \quad \text{(S7)}$$

Multiplying by $\frac{S_m}{T_{leaf}} V_{strom}$ converts carboxylation from per area to per stroma volume units. $k_c$ is the catalytic rate of Rubisco ($m^{-1}$) and $K_m$ is effective Michaelis-Menten constant for Rubisco (mol $m^{-3}$). Following Earles et al. (2017), we assumed the relative concentration of Rubisco follows that of Nishio, Sun, and Vogelmann (1993), but scaled such that the bulk leaf Rubisco concentration integrates to $X_c$ described in Table S1. We estimated a continuous function describing the relative Rubisco profile as a function of leaf depth using a generalized additive model with the `gam()` function in *R* package **mgcv** version 1.9.0 (Wood 2017).

The effective photosynthetic $e^-$ transport rate ($j_e$) is the minimum of the maximum ($j_{max}$) and potential ($j_\infty$) photosynthetic $e^-$ transport rates at each position within the mesophyll:

$$j_e = \min(j_{\text{max}}, j_\infty). \tag{S8}$$

The local $j_{\text{max}}$ follows the same depth profile as Rubisco and is scaled by local $S_m$ and $V_{\text{strom}}$ so that it integrates to $J_{\text{max}}$ on a leaf-area basis (Earles et al. 2017):

$$J_{\text{max}} = \int_0^{T_{\text{leaf}}} j_{\text{max},i} S_{\text{m},i} V_{\text{strom}} dz. \tag{S9}$$

Potential $e^-$ transport follows the same depth profile as chlorophyll concentration and is scaled by local $S_m$ and $V_{\text{strom}}$ so that it integrates to $J_\infty$ on a leaf-area basis (Earles et al. 2017), where:

$$J_\infty = I_0 \alpha \beta \phi_{\text{PSII}} = \int_0^{T_{\text{leaf}}} j_{\infty,i} S_{\text{m},i} V_{\text{strom}} dz. \tag{S10}$$

Potential $e^-$ transport is a product of PPFD incident on the leaf surface ($I_0$, mol $m^{-2}$ $s^{-1}$), whole-leaf light absorption ($\alpha$, mol $mol^{-1}$), the fraction of light absorbed by PSII ($\beta$, mol $mol^{-1}$), and the quantum yield of PSII $e^-$ transport ($\phi_{\text{PSII}}$, mol $mol^{-1}$).

The local chlorophyll concentration is a function of depth described by a quadratic equation (Johnson et al. 2005):

$$F_{\text{chl}} = f(z) = b_{0,\text{chl}} + b_{1,\text{chl}} z + b_{2,\text{chl}} z^2. \tag{S11}$$

We used generic parameters from Borsuk and Brodersen (2019), rescaled to relative depth on a 0-1 rather than 0-100 interval.

For mass balance, the $CO_2$ assimilated via carboxylation must be supplied by diffusive flux from the intercellular airspace into the chloroplast stroma. The volumetric rate of $CO_2$ diffusion from the intercellular airspace into the chloroplast stroma, $f_{\text{liq}}$, is:

$$f_{\text{liq}} = \frac{g_{\text{liq}}(C_{\text{liq}} - C_{\text{ias}})}{T_{\text{leaf}}/S_m}, \tag{S12}$$

where $g_{\text{liq}}$ is the $CO_2$ conductance from the intercellular airspace into the chloroplast stroma (m $s^{-1}$), $C_{\text{liq}}$ (mol $m^{-3}$) is the [$CO_2$] in the stroma, and $S_m$ is mesophyll surface area-to-leaf surface area ratio. Noting that $g_{\text{liq}}$ is conductance per $m^2$ of stroma, this means the length scale to divide by should be the inverse of stroma area per unit bulk leaf volume, i.e. $1/[S_c(1/T_{\text{leaf}})] = T_{\text{leaf}}/S_c$. For simplicity, we assume that the entire mesophyll surface area is lined with chloroplasts, hence $S_m = S_c$. Following Earles et al. (2017), we assumed greater surface area in the palisade ($S_{\text{m,pal}}$) than spongy ($S_{\text{m,spg}}$) mesophyll. The fractions of palisade and spongy mesophyll are $f_{\text{pal}}$ and $f_{\text{spg}}$, respectively.

Following Earles et al. (2017), we set boundary conditions at the stomata, $C_{\text{stom}}$, to be $0.85\times$ atmospheric $CO_2$ (Table S1). The fluxes on the left and right sides are 0 because of symmetry; the other fluxes on the ab- and adaxial surface are assumed 0 (i.e. the epidermis is impermeable to $CO_2$).

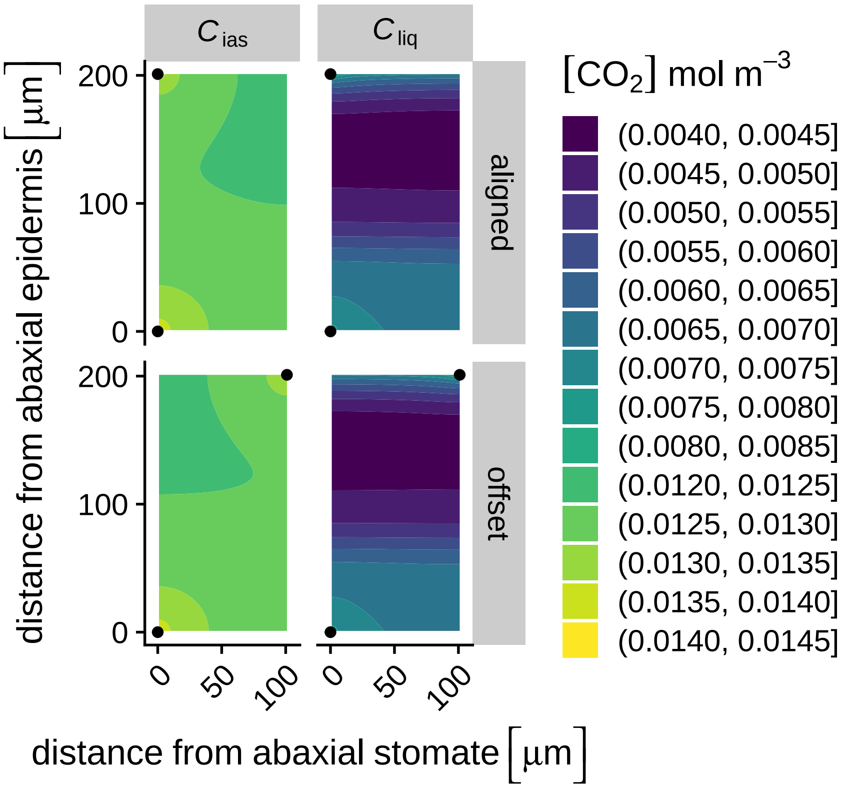


Figure S1: Example profiles of volumetric $CO_2$ concentrations within otherwise identical amphistomatous leaves that have stomatal positions offset (top row) or aligned (bottom row) based on the 2-D porous medium model. Stomatal positions are indicated by black points at the top and bottom of panels. When stomata are aligned, both ab- and adaxial stomata are position 0 along the $x$-axis; when stomata are offset, the adaxial stomate is positioned $U/2$ distance away. In this example, variables are set as: $I_0 = 1000$ $\mu$mol m$^{-2}$ s$^{-1}$; $\varphi_{\text{pal}} = 0.2$ m$^3$ airspace m$^{-3}$ leaf; $T_{\text{leaf}} = 200$ $\mu$m; $U = 200$ $\mu$m. All other parameter values are described in Table S1. $C_{\text{ias}}$ = [$CO_2$] in intercellular airspace; $C_{\text{liq}}$ = [$CO_2$] in chloroplast stroma; $I_0$ = PPFD incident on the leaf surface; $\varphi_{\text{pal}}$ = Fraction of intercellular airspace (aka porosity), palisade; $T_{\text{leaf}}$ = Leaf thickness; $U$ = Interstomatal distance.

Table S1: Glossary of model terms and mathematical symbols.

| Name | Symbol | Value | Units | Notes |
|---|---|---|---|---|
| Whole-leaf light absorption | $\alpha$ | 0.8 | mol mol$^{-1}$ | assumed |
| Chlorophyll spatial distribution coefficient | $b_{0,\text{chl}}$ | 67.5 | — | Borsuk and Brodersen (2019); Equation S11 |
| Chlorophyll spatial distribution coefficient | $b_{1,\text{chl}}$ | 41.5 | — | Borsuk and Brodersen (2019); Equation S11 |
| Chlorophyll spatial distribution coefficient | $b_{2,\text{chl}}$ | −29 | — | Borsuk and Brodersen (2019); Equation S11 |
| Fraction of light absorbed by PSII | $\beta$ | 0.5 | mol mol$^{-1}$ | assumed |
| [$CO_2$] in intercellular airspace | $C_{\text{ias}}$ | calculated | mol m$^{-3}$ | Equation S1 and Equation S12 |
| [$CO_2$] in chloroplast stroma | $C_{\text{liq}}$ | calculated | mol m$^{-3}$ | Equation S1 |
| [$CO_2$] in substomatal cavity | $C_{\text{stom}}$ | $1.50 \times 10^{-2}$ | mol m$^{-3}$ leaf | assumed |
| [$CO_2$] compensation point | $\Gamma^*$ | $1.35 \times 10^{-3}$ | mol m$^{-3}$ stroma | Caemmerer (2000) |
| Diffusivity of [$CO_2$] in intercellular airspace | $D_{\text{c}}$ | $1.54 \times 10^{-5}$ | m$^2$ s$^{-1}$ | assumed |
| Effective diffusivity of [$CO_2$] in intercellular airspace | $D_{\text{e}}$ | calculated | m$^2$ s$^{-1}$ | Equation S3 |
| Fraction of palisade mesophyll | $f_{\text{pal}}$ | 0.6 | 1 | $1 = f_{\text{pal}} + f_{\text{spg}}$ |
| Fraction of spongy mesophyll | $f_{\text{spg}}$ | 0.4 | 1 | $1 = f_{\text{pal}} + f_{\text{spg}}$ |
| Chlorophyll fluoresence profile along leaf depth normalized by total fluoresence | $F_{\text{chl}}$ | calculated | 1 | Borsuk and Brodersen (2019); Equation S11 |
| Conductance of cell wall, plasmalemma, cytosol, chloroplast envelope, and chloroplast stroma | $g_{\text{liq}}$ | $2.50 \times 10^{-4}$ | m$^3$ m$^{-2}$ stroma s$^{-1}$ | Evans et al. (2009) |
| PPFD incident on the leaf surface | $I_0$ | variable | mol m$^{-2}$ s$^{-1}$ | assumed |
| Potential photosynthetic e$^-$ transport rate on a leaf area basis | $J_\infty$ | calculated | mol m$^{-2}$ leaf s$^{-1}$ | Equation S10 |
| Maximum photosynthetic e$^-$ transport rate on a leaf area basis | $J_{\text{max}}$ | $2.75 \times 10^{-4}$ | mol m$^{-2}$ leaf s$^{-1}$ | assumed |

| Name | Symbol | Value | Units | Notes |
|---|---|---|---|---|
| Effective photosynthetic $e^-$ transport rate on a stroma volume basis | $j_e$ | calculated | mol $m^{-3}$ stroma $s^{-1}$ | Equation S8 |
| Potential photosynthetic $e^-$ transport rate on a stroma volume basis | $j_\infty$ | calculated | mol $m^{-3}$ stroma $s^{-1}$ | Equation S10 |
| Maximum photosynthetic $e^-$ transport rate on a stroma volume basis | $j_{max}$ | calculated | mol $m^{-3}$ stroma $s^{-1}$ | Equation S9 |
| Catalytic rate of Rubisco | $k_c$ | 2.84 | $m^{-1}$ | Tholen and Zhu (2011) |
| Rubisco effective $K_m$ | $K_m$ | $1.87 \times 10^{-2}$ | mol $m^{-3}$ | Caemmerer (2000) |
| Number of elements in $x$ direction | $n_x$ | calculated | — | $U = 2n_x t_{elem}$ |
| Number of elements in $z$ direction | $n_z$ | calculated | — | $T_{leaf} = n_z t_{elem}$ |
| Fraction of intercellular airspace (aka porosity), palisade | $\varphi_{pal}$ | variable | $m^3$ airspace $m^{-3}$ leaf | assumed |
| Fraction of intercellular airspace (aka porosity), spongy | $\varphi_{spg}$ | 0.3 | $m^3$ airspace $m^{-3}$ leaf | assumed |
| Quantum yield of PSII $e^-$ transport | $\phi_{PSII}$ | 0.85 | mol $mol^{-1}$ | assumed |
| Volumetric rate of RuBP carboxylation | $r_c$ | calculated | mol $m^{-2}$ stroma $s^{-1}$ | Equation S5 |
| Volumetric respiration rate | $r_d$ | $6.60 \times 10^{-2}$ | mol $m^{-2}$ stroma $s^{-1}$ | Earles et al. (2017); Tholen and Zhu (2011) |
| Volumetric rate of photorespiratory $CO_2$ release | $r_p$ | calculated | mol $m^{-2}$ stroma $s^{-1}$ | Earles et al. (2017); Equation S4 |
| Mesophyll surface area-to-leaf surface area ratio, palisiade | $S_{m,pal}$ | 20 | $m^2$ mesophyll $m^{-2}$ leaf | assumed |
| Mesophyll surface area-to-leaf surface area ratio, spongy | $S_{m,spg}$ | 2 | $m^2$ mesophyll $m^{-2}$ leaf | assumed |
| Tortuosity of intercellular airspace | $\tau$ | 1.55 | m $m^{-1}$ | Syvertsen et al. (1995) |
| Thickness of element in both $x$ and $z$ directions | $t_{elem}$ | $1.00 \times 10^{-6}$ | m | $T_{leaf} = n_z t_{elem}$ |

| Name | Symbol | Value | Units | Notes |
|---|---|---|---|---|
| Leaf thickness | $T_{\text{leaf}}$ | variable | m | $T_{\text{leaf}} = n_z t_{\text{elem}}$ |
| Interstomatal distance | $U$ | variable | m | $U = n_x t_{\text{elem}}$ |
| Stroma volume-to-mesophyll surface area ratio | $V_{\text{strom}}$ | $1.74 \times 10^{-6}$ | $m^3$ stroma $m^{-2}$ mesophyll | Earles et al. (2017); Tholen and Zhu (2011) |
| Rubisco-limited carboxylation rate | $w_{\text{c}}$ | calculated | mol $m^{-2}$ stroma $s^{-1}$ | Equation S6 |
| RuBP regeneration-limited carboxylation rate | $w_{\text{j}}$ | calculated | mol $m^{-2}$ stroma $s^{-1}$ | Equation S7 |
| Rubisco concentration in stroma | $X_{\text{c}}$ | 2.5 | mol $m^{-3}$ stroma | Tholen and Zhu (2011); Oguchi, Hikosaka, and Hirose (2003) |
